# Split-Small GTPase Reassembly as a Method to Control Cellular Signaling with User-Defined Inputs

**DOI:** 10.1101/2025.01.28.635345

**Authors:** Yuchen He, Benjamin M. Faulkner, Emily Hyun, Cliff I. Stains

**Affiliations:** Department of Chemistry, University of Virginia, Charlottesville, VA 22904, USA; University of Virginia Cancer Center, University of Virginia, Charlottesville, VA 22908, USA; Virginia Drug Discovery Consortium, Blacksburg, VA 24061, USA

## Abstract

Small GTPases are critical signaling enzymes that control diverse cellular functions such as cell migration and proliferation. However, dissecting the roles of these enzymes in cellular signaling is hindered by the lack of a plug-and-play methodology for the direct, temporal control of small GTPase activity using user-defined inputs. Herein, we present a method that pairs split-GTPases with user-defined chemical inducer of dimerization (CID) systems in a plug-and-play manner to directly control small GTPase signaling in living cells. The modularity of split-small GTPase systems allows for the selection of CIDs with minimal off-target effects on the pathway being studied. Our results highlight the ability to obtain consistent pathway activation with varying CID systems for direct control of MAPK signaling, filopodia formation, and cell retraction. Thus, split-small GTPase systems provide a customizable platform for development of temporally gated systems for directly controlling cellular signaling with user-defined inputs.

## Introduction

Small GTPases are a family of hydrolase enzymes that play a crucial role in regulating diverse cellular processes including cell growth and proliferation as well as cell motility.^1-3^ These enzymes act as molecular switches, alternating between an active GTP-bound state and an inactive GDP-bound state.^1^ The activation of these enzymes is typically facilitated by guanine nucleotide exchange factors (GEFs), while deactivation is promoted by GTPase-activating proteins (GAPs).^4^ Mutations that influence the activation state of small GTPases are associated with a variety of diseases, such as cancer.^5-7^ Due to their pivotal role in fundamental cellular signaling pathways and their implication in a variety of diseases, methods to control the activity of these enzymes are highly desirable for both biomedical research as well as synthetic biology applications.

Approaches for assessing the involvement of small GTPases in cellular signaling include genetic manipulation,^8,9^ biochemical characterization,^10-12^ and live-cell imaging techniques.^13,14^ While these methods have contributed to important advances in the understanding of small GTPase signaling, they are not without limitations. For example, genetic manipulation, such as the overexpression of mutant GTPases in living cells, may induce compensatory signaling changes that obscure the function of the target GTPase.^15,16^ Biochemical characterization methods, such as pull-down assays, capture specific GTPase interactions but may not fully reflect the range of activity states or context-dependent regulatory interactions.^17^ Imaging-based techniques are valuable for studying dynamic processes but often require complex setups to obtain quantitative data, limiting their accessibility. To address these issues, protein engineering-based approaches have been developed that enable temporal control of small GTPase activity within living cells.^18-20^ Although these approaches represent valuable tools, they often require extensive, case-by-case optimization for each new small GTPase target. Moreover, the ability to control small GTPase activity with user-defined inputs is challenging, necessitating reliance on small molecule inputs with potential off-target effects on the pathway being studied. Given the importance of small GTPase signaling in both normal and disease-relevant processes,^5-7^ there is a critical need for the development of a plug-and-play method to control the activity of a target small GTPase with user-defined inputs.

To address the first technical hurdle for realization of such an approach, our lab has recently disclosed the development of a generalizable system for temporally controlling the activity of a target small GTPase.^21^ Specifically, we showed that a fragmentation site termed N12/13C, discovered in Cdc42,^22^ could be applied across the small GTPase superfamily using sequence alignment (**Figure S1**), yielding functional, split-small GTPases without the need for case-by-case optimization. More specifically, utilizing the rapamycin-dependent association of FKBP and FRB, we demonstrated the ability to temporally control the activity of KRas, Cdc42, and RhoA (**Figure 1a** and **b**).^21^ However, rapamycin is a well-known mTOR kinase inhibitor,^23-25^ and we have shown that its off-target effects can obscure the activity of split-small GTPases (such as split-KRas) within living cells.^21^ In a broader sense, the ability to control split-small GTPase activity with user-defined inputs would offer a versatile system for applications in synthetic biology as well as biomedical research. Herein, we demonstrate the ability to control split-GTPase function in living cells with user-defined inputs such as abscisic acid (ABA) or gibberellic acid (GA) (**Figure 1c**).^26-28^ We demonstrate that comparable split-GTPase activation in living cells can be achieved using these different CID systems. These data clearly demonstrate the ability to employ user-defined inputs to activate small GTPase signaling in living cells and provide a generalizable, plug-and-play system for the temporal control of small GTPase signaling.

**Figure 1.**
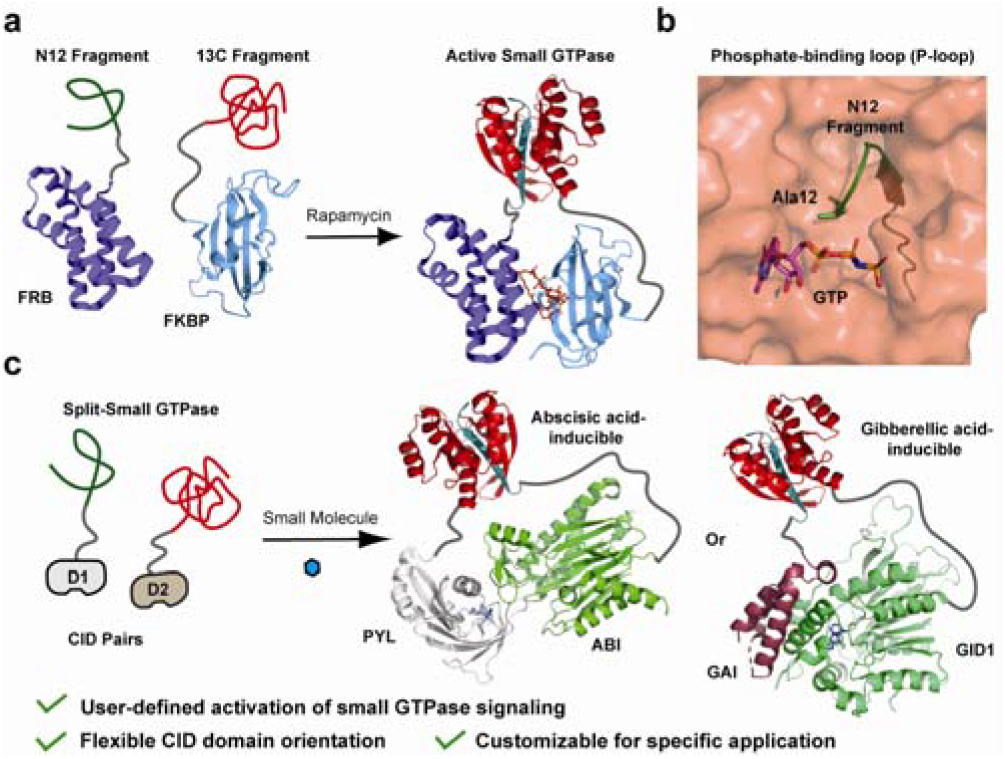
User-defined inputs for controlling split-small GTPase reassembly. **a)** A schematic of our previous work in which rapamycin-induced reassembly of a split-small GTPase was used to temporally control cell signaling (structures used were PDB: 6M4U and 1E0A). **b)** The crystal structure of active Cdc42 (PDB: 1E0A) is shown with the N12 fragmentation site in the phosphate-binding loop (P-loop) highlighted in green. Fragmentation of the enzyme at this position leads to production of inactive fragments that can be reconstituted using concentration-induced reassembly. **c)** Concentration-induced reassembly of split-small GTPase allows user-defined inputs for reassembly to be employed. For example, the rapamycin dependent CID can be replaced with an abscisic acid- or gibberellic acid-based system depending on experimental needs (PDB: 3KDJ and 2ZSH).

## Results and Discussion

### Split-GTPase construct designs

To investigate the plug-and-play nature of our split-small GTPase reassembly systems, we envisioned replacing rapamycin-based CIDs used previously.^21^ These new constructs consist of fusions between either the N12 or 13C small GTPase fragments, the appropriate CID domain, and a fluorescent protein. In each case, fusions are made at the native termini of the N12 and 13C to allow for productive reassembly. A CAAX motif derived from KRas4b^29^ is utilized to localize the 13C fragment to the cytosolic face of the cell membrane, mirroring the native localization of small GTPases (**Figure 2a**).^30^ In the presence of a small molecule input, dimerization of the CID domains leads to an increase in the local concentration of small GTPase fragments, resulting in reassembly and activation of signaling.^21,22^ Fluorescent proteins, mCerulean and mVenus, are incorporated to monitor relative expression levels of each fragment and to confirm appropriate localization. Both protein fusions are expressed from a single vector, pIRES, which features an internal ribosome entry site (IRES) (**Figure 2a**).

**Figure 2.**
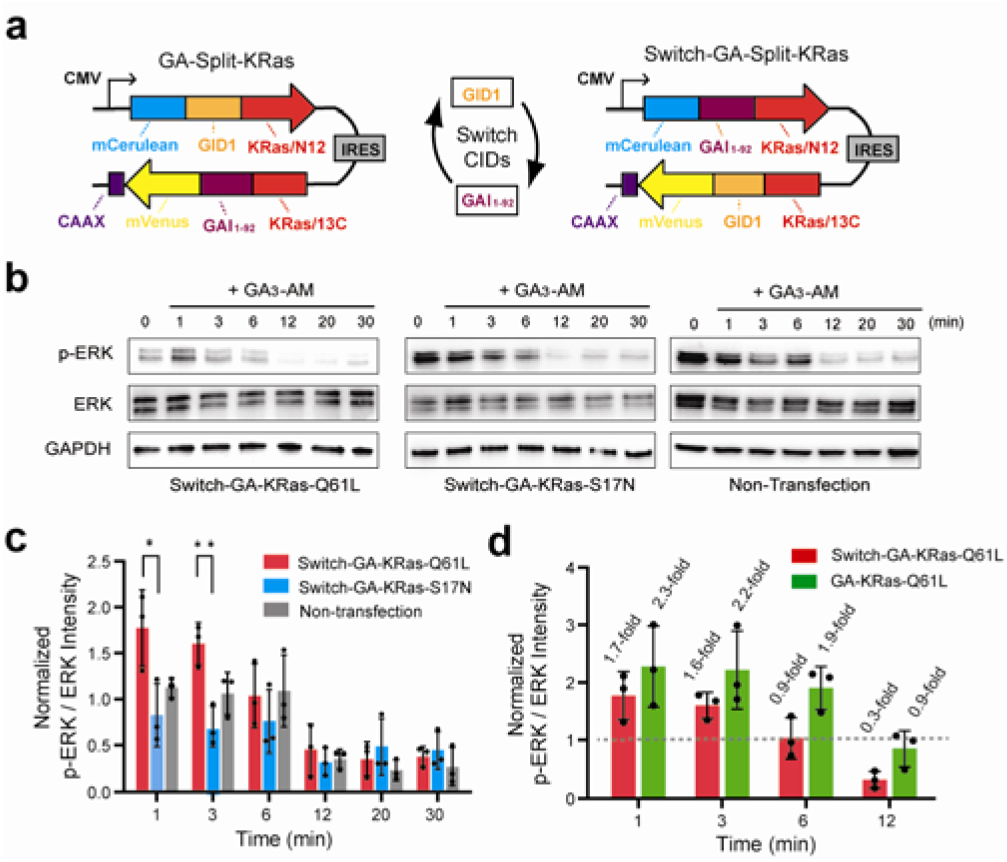
The influence of CID domain orientation on split-KRas signaling. **a)** Design of constructs with switched orientations of GA-gated CID domains for controlling split-KRas protein reassembly. **b)** Representative western blots from HeLa cells transfected with Switch-GA-Split-KRas constructs and stimulated with 10 µM gibberellin (GA_3_-AM) for the indicated time. **c)** Quantified band intensities from panel **b** for p-ERK relative to total ERK from three independent biological replicates (mean ± SD). * is adjusted P<0.05 and ** is adjusted P<0.01 from one-way ANOVA followed by a Fisher’s PLSD test. **d)** Comparison of ERK activation for the constructs shown in panel **a**.

### The effect of CID domain orientation on gibberellic acid-induced split-KRas signaling in living cells

We have previously disclosed a split-KRas construct which relies on GA-gated association of GID1 and GAI^26^ to activate KRas signaling in HeLa cells.^21^ In HeLa cells expressing constitutively active GA-KRas-Q61L fragments, we observed a clear 2.3-fold increase in ERK phosphorylation one minute after stimulation with GA_3_-AM, which returned to baseline within 12 minutes.^21^ This result highlights the ability to activate split-KRas using GA as an input. However, previous work has shown that the efficiency of split-protein reassembly can vary depending on CID domain orientation.^31^ To address the generality of the split-KRas system, we asked whether the orientation of the CID domains influence KRas signaling as assessed by ERK phosphorylation. Accordingly, we constructed a switched version of the GA-gated split-KRas system, in which the orientation of GAI and GID1 are swapped (**Figure 2a**). After transiently transfecting HeLa cells with this new construct, we confirmed the localization of the split-KRas fragments using confocal microscopy (**Figure S2a**) and equivalent transfection efficiency of constitutively active (Q61L) and dominant negative (S17N) Switch-GA-Split-KRas constructs (**Figure S2b**). To assess the activation of Switch-GA-Split-KRas constructs, cells were stimulated with 10 µM GA_3_-AM and the phosphorylation status of ERK was probed via Western blotting. These experiments demonstrated a reproducible 1.7-fold increase in ERK phosphorylation that returned to baseline by 6 minutes. (**Figure 2b, 2c**, and **Figure S2c**). In contrast, cells expressing Switch-GA-KRas-S17N or non-transfected cells showed no significant changes in ERK phosphorylation (**Figure 2c**). When compared to our previous GA-Split-KRas results,^21^ a similar magnitude of ERK phosphorylation is observed with different orientations of CID proteins in the split-KRas system (**Figure 2d**). However, the GA-Split-KRas construct appears to produce more prolonged activation of ERK (e.g. 6 min time point, **Figure 2d**). These results underscore the modular nature of split-GTPase fragments. Although we cannot fully rule out context-dependent effects on reassembly efficiency for all CID domains, these results imply that comparable reassembly efficiency can be achieved when the CID domain termini are on the same face of the CID complex (**Figure S3**). Interestingly, we observed a decrease in average ERK phosphorylation in cells expressing the Switch-GA-KRas-S17N construct (**Figure 2c**). We hypothesize that the reassembly of dominant negative split-KRas fragments may act as competitive inhibitors of upstream GEFs, leading to reduced pathway activation. Our lab is currently investigating the use of split-dominant negative small GTPases as inhibitors of cellular signaling. Overall, these results reinforce the modular nature of split-small GTPases.

### Controlling filopodia formation using GA as an input

We have previously demonstrated rapamycin-based control of filopodia formation in living cells using split-Cdc42.^21^ Although this approach enabled temporal control of filopodia formation, off-target effects from rapamycin could complicate the analysis of downstream effectors activated by Cdc42. To address this issue, we explored the use of alternative, user-defined inputs to control split-Cdc42 reassembly in living cells. Accordingly, we replaced the rapamycin binding domains in our original system with the GA-dependent domains GID1 and GAI (**Figure 3a**) and confirmed the localization of each fragment following transient transfection (**Figure S4**). Subsequently, cells were stimulated with 10 µM GA_3_-AM and filopodia formation was monitored using confocal microscopy. Gratifyingly, we detected a distinct increase in filopodia formation after GA_3_-AM stimulation in HeLa cells expressing GA-Split-Cdc42-Q61L (**Figure 3b**, arrows). In contrast, cells expressing GA-Split-Cdc42-T17N displayed no observable change in membrane morphology (**Figure 3b**). To assess the reproducibility of this GA-gated system, we employed FiloQuant to quantify filopodia formation across multiple transfected cells.^32^ In this analysis, increased filopodia formation was only observed in GA-Split-Cdc42-Q61L expressing cells treated with GA_3_-AM (**Figure 3c**). On average this system showed a reduced fold increase in filopodia formation (1.6-fold) compared to the rapamycin system (4.1-fold, **Figure 3d**), potentially due to the weaker binding affinity of GA_3_-AM with GID1 and GAI compared to that of rapamycin with FKBP and FRB.^33,34^ Notably, filopodia formation induced by GA_3_-AM occurred on a faster timescale (∼15 min) than with the rapamycin-gated system (∼30 minutes, **Figure S5**). Thus, while the GA system produces a smaller fold increase in filopodia formation on average, its enhanced temporal control could be beneficial for studying cell signaling pathways on a more rapid timescale. Furthermore, GA is less likely to produce off-target effects in mammalian cells compared to rapamycin. Similar to our previous rapamycin-based split-Cdc42 system,^21^ we only observed long finger-like filopodia (Cdc42 phenotype) as opposed to broader lamellipodia structures (Rac1 phenotype) with the GA-activated system (**Figure 3b**). Previous approaches to modulate upstream regulators of Cdc42 have often resulted in both Cdc42 and Rac1 phenotypes due to cross-reactivity of regulatory proteins.^20,35,36^ Thus, this work further reinforces the benefits of directly activating small GTPases with user-defined inputs.

**Figure 3.**
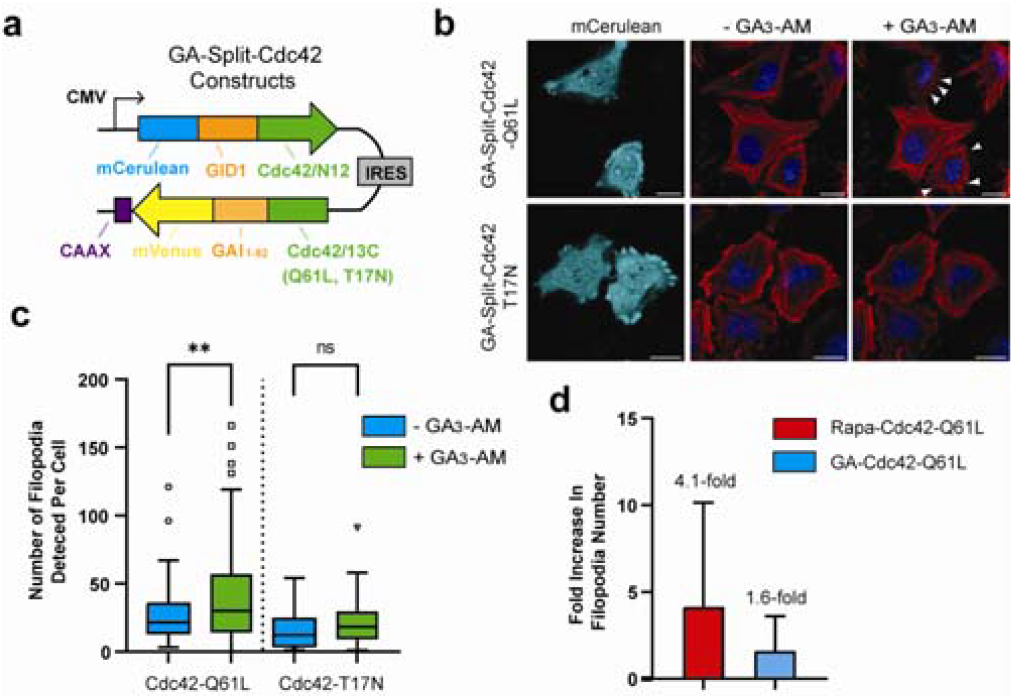
GA-gated Cdc42 is capable of regulating filopodia formation in mammalian cells. **a)** A construct for dual expression of GA-Split-Cdc42 proteins. **b)** Confocal images of HeLa cells in the mCerulean channel (representing transfected cells) or merged images of CellMask Deep Red Actin tracker (red) and Hoechst stain (blue) in the absence or presence of 10 µM GA_3_-AM for 15 min. Cells expressing GA-Split-Cdc42-Q61L display clear filopodia formation upon GA_3_-AM stimulation (arrows). **c)** Quantified filopodia number from panel **(b)** for n > 50 cells using FiloQuant shows a 1.6-fold increase in the number of filopodia formed in GA_3_-AM treated cells expressing split-Cdc42-Q61L. **d)** Comparison of filopodia formation in rapamycin-gated and GA-gated split-Cdc42 systems in HeLa cells. Scale bar represents 20 µm.

### Controlling cell retraction using abscisic acid as an input

We next investigated whether the reassembly of split-RhoA and subsequent cell retraction could be controlled using ABA-mediated dimerization of PYL and ABI.^27^ Thus, we constructed ABA-gated split-RhoA constructs (**Figure 4a**) and transiently transfected them into HeLa cells. Confocal imaging confirmed appropriate localization of the fragmented RhoA proteins (**Figure S6a**). Notably, HeLa cells expressing ABA-Split-RhoA-Q63L displayed significant membrane retraction following ABA stimulation (**Figure 4b**, white arrows). Non-transfected cells within the same field of view served as internal controls and showed no response to ABA. Cells expressing dominant negative ABA-Split-RhoA-T19N did not display substantial cell retraction upon stimulation (**Figure S6b**).

**Figure 4.**
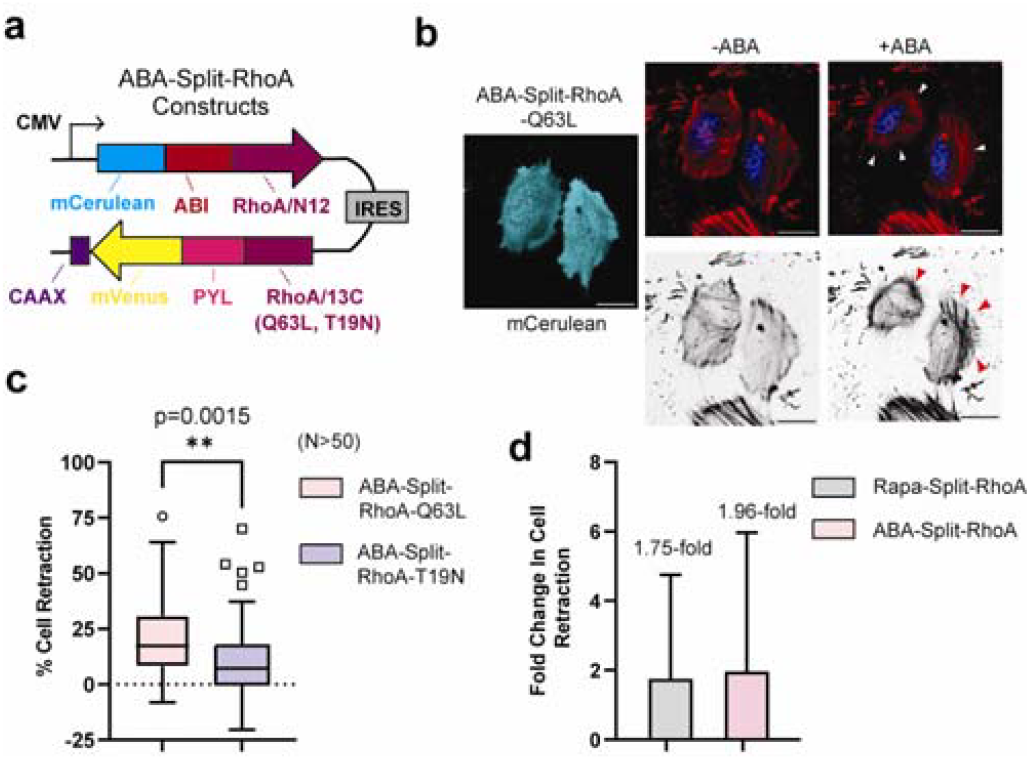
ABA-gated RhoA can control cell retraction in HeLa cells. **a)** Design of dual expression constructs for ABA-Split-RhoA proteins. **b)** Confocal images of HeLa cells in the mCerulean channel (representing transfected cells) or merged images of CellMask Deep Red Actin tracker (red) and Hoechst stain (blue) in the absence or presence of 100 µM ABA for 30 min. Cells expressing ABA-Split-RhoA-Q63L display clear membrane retraction upon ABA stimulation (white arrows). Formation of contractile F-actin filaments is also observed post stimulation (red arrows). **c)** Quantification of transfected cells from panel **(b)** showing the percent retraction of each cell for ABA-Split-RhoA-Q63L and ABA-Split-RhoA-T19N systems after treatment with ABA for 30 min. **d)** Comparison of cell retraction for rapamycin-gated and ABA-gated split-RhoA systems in HeLa cells. Scale bar represents 20 µm.

Additionally, in ABA-Split-RhoA-Q63L expressing cells, ABA treatment resulted in elongated, contractile F-actin filaments converging at the nucleus (**Figure 4b**, red arrows). To quantify the effects of ABA on RhoA-mediated cell retraction, we employed ImageJ to measure the area of each transfected cell before and after ABA treatment. Results revealed a clear increase in cell retraction for the constitutively active mutant (ABA-Split-RhoA-Q63L) compared to the dominant negative mutant (ABA-Split-RhoA-T19N) after treatment with ABA (**Figure 4c**). Furthermore, this ABA-gated system resulted in a 1.96-fold increase in cell retraction, which is comparable to the 1.75-fold change observed with our previously described rapamycin-gated split-RhoA system (**Figure 4d**). These results again underscore the ability to generate split-GTPase systems capable of responding to user-defined inputs.

## Conclusions

In summary, we have demonstrated that split-small GTPases can be combined with user-defined inputs to enable temporal control of signaling pathways within living cells. The orientation of the domains used to trigger split-small GTPase reassembly did not influence the magnitude of signaling (**Figure 2d**), although we note that the termini of CID domains used here are on the same face of the complex (**Figure S3**). We further demonstrated the ability to temporally control diverse cellular functions such as filopodia formation (**Figure 3**) and cell retraction (**Figure 4**), through the use of different small molecule inputs. Importantly, the reassembly of split-small GTPase enables direct activation of a given small GTPase, avoiding cross-talk from upstream regulatory proteins. Given the potential off-target effects of small molecules on cellular signaling, the ability to tailor split-small GTPase reassembly to fit the needs of an experiment represents an important feature for biomedical research and synthetic biology applications. Since rapamycin-, gibberellic acid-, and abscisic acid-based systems are orthogonal, this work opens up the possibility for orthogonal activation of multiple split-small GTPases within living cells. Ultimately, we envision the plug-and-play nature of split-small GTPases will enable the development of user-defined systems for uncovering the roles of these enzymes in cell signaling.

## Acknowledgements

We thank the W.M. Keck Center for Cellular Imaging for the use of the Leica STELLARIS 8 confocal/FLIM/tauSTED microscope system (NIH OD030409) and members of the C. Stains lab for editing the manuscript and helpful conversations. We acknowledge financial support from the NIH (R35GM148221) and the University of Virginia. The content of this work is solely the responsibility of the authors and does not necessarily represent the official views of the NIH.

## Table of Contents Figure

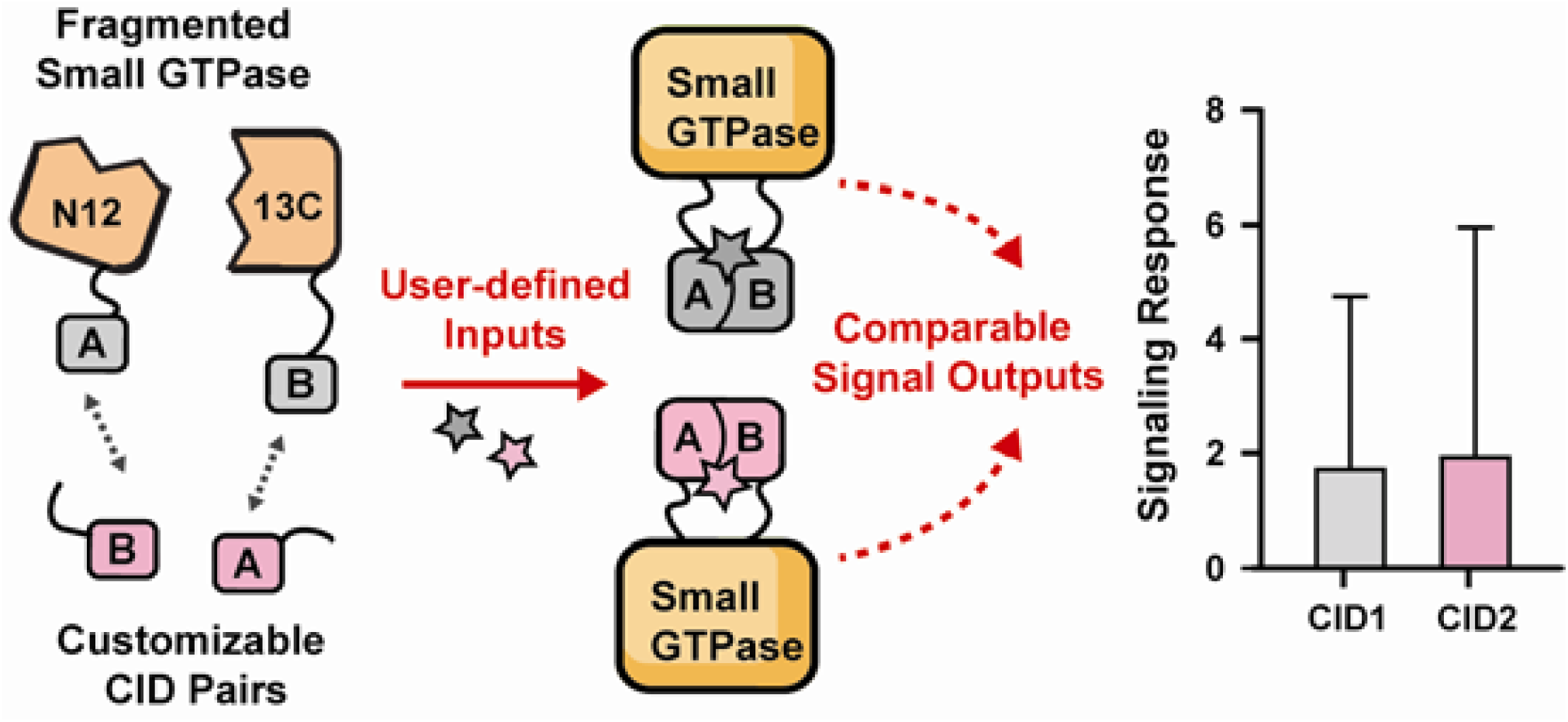

