## Supplementary Material for "Split-Small GTPase Reassembly as a Method to Control Cellular Signaling with User-Defined Inputs"

**Figure S1**

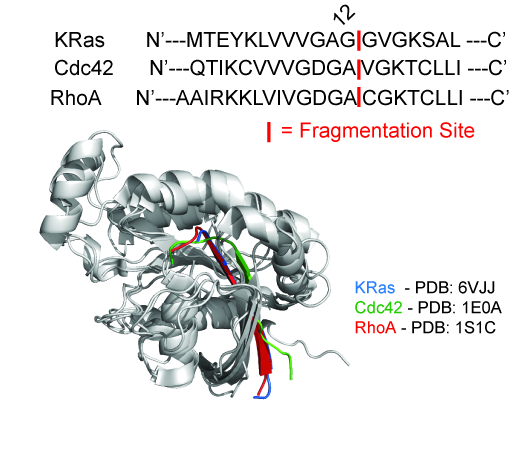

**Figure S1**. Application of the N12/13C fragmentation site across the small GTPase superfamily. A sequence alignment for KRas, Cdc42, and RhoA showing the N12/13C fragmentation site is shown (top). Numbering refers to Cdc42. An overlay of crystal structures corresponding to the indicated, active small GTPase are shown (bottom). The N12 fragment of each small GTPase is highlighted.

**Figure S2**

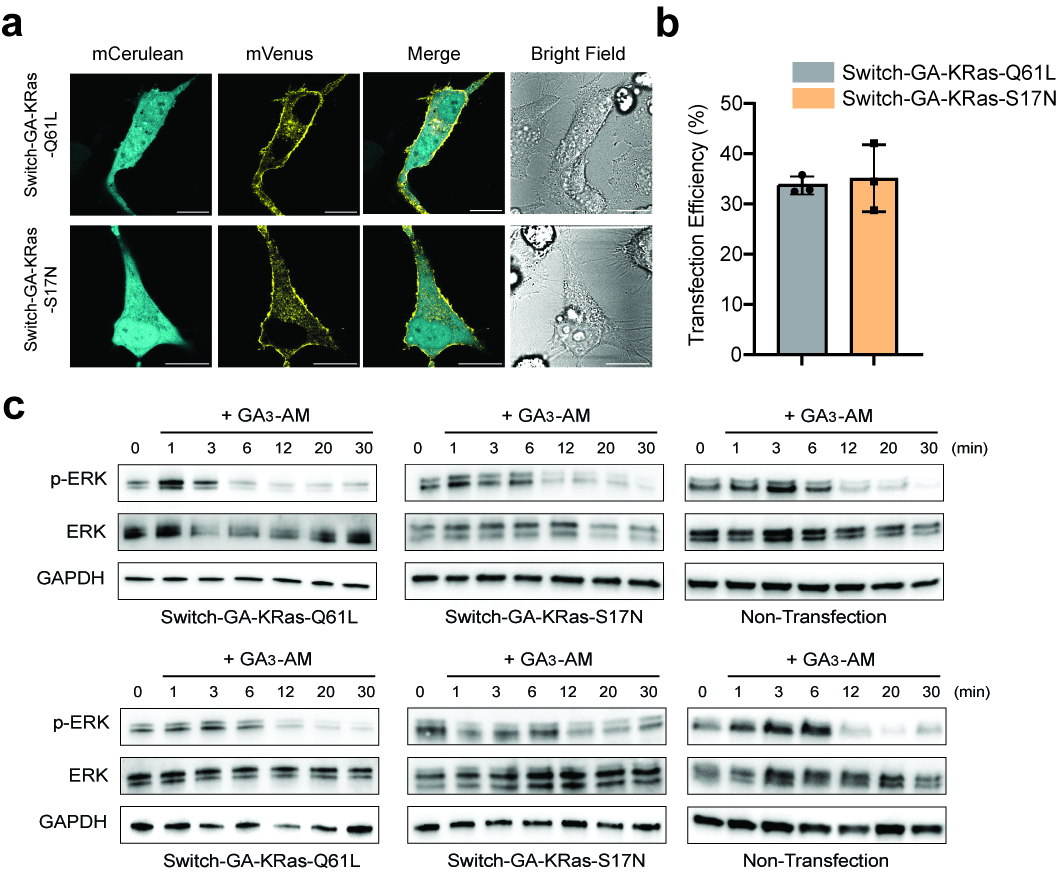

**Figure S2**. Switch-GA-Split-KRas activates ERK signaling. **a)** Confocal images of HeLa cells expressing Switch-GA-KRas-Q61L or Switch-GA-KRas-S17N constructs. mCerulean fluorescence is observed in the cytosol while the mVenus fluorescence is localized to the cell membrane. Scale bar represents 20 µm. **b)** Transfection efficiency of Switch-GA-Split-KRas constructs in HeLa cells was determined by normalizing the number of fluorescent cells to the total observed cells in the bright field channel. Data points are biological replicates in which > 50 cells are imaged in each replicate. **c)** Western blots for biological replicates assessing phosphorylation of ERK using Switch-GA-Split-KRas. HeLa cells were stimulated with 10 µM gibberellin (GA_3_-AM) for the indicated time, lysed, and Westerns were performed with the indicated antibody.

**Figure S3**

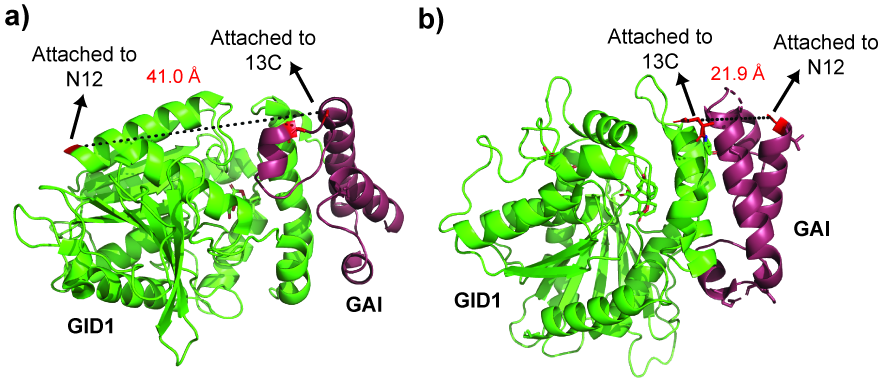

**Figure S3.** The termini of the GAI and GID1 are on the same face of the GA-mediated complex (PDB: 2ZSH). **a)** GAI and GID1 termini used for fusion in the GA-KRas-Q61L construct. **b)** GAI and GID1 termini used for fusion in the Switch-GA-KRas-Q61L construct. Fusion of these termini to either N12 or 13C using flexible 17-30 amino acid linkers does not appreciably influence reassembly (see **Figure 2**).

**Figure S4**

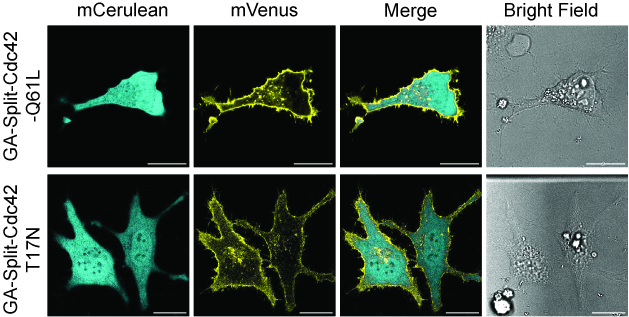

**Figure S4.** Confocal images of HeLa cell expressing the GA-Split-Cdc42 fragments. mCerulean fluorescence is observed in the cytosol while the mVenus fluorescence is localized to the cell membrane for both GA-Split-Cdc42-Q61L and GA-Split-Cdc42-T17N.

**Figure S5**

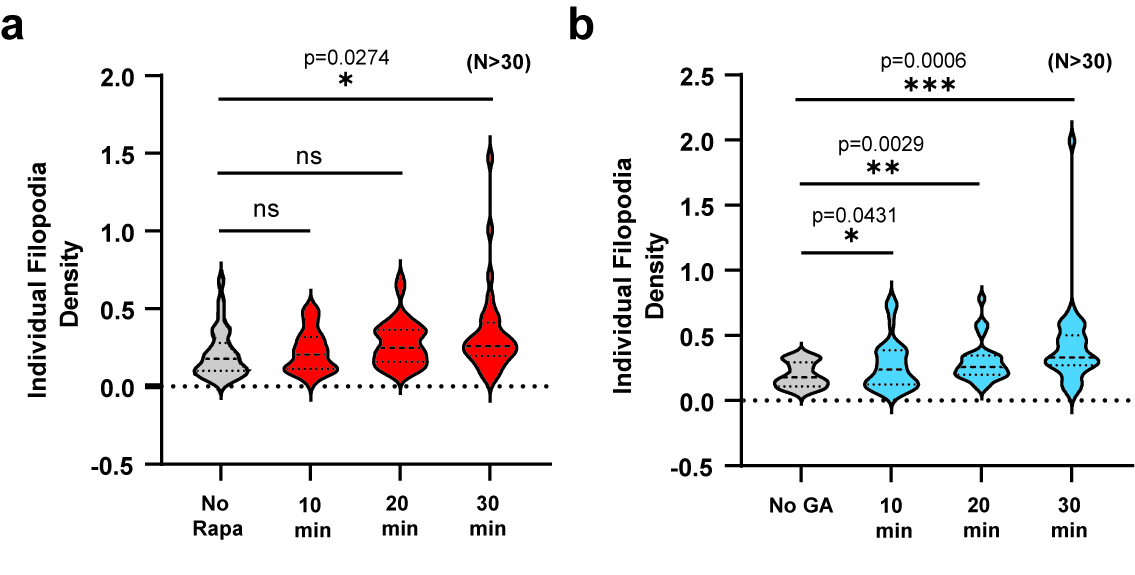

**Figure S5**. Time-course comparison of rapamycin-gated and gibberellic acid-gated filopodia formation using split-Cdc42-Q61L. **a)** Filopodia density in HeLa cells transfected with Rapa-Split-Cdc42-Q61L was measured at the indicated time points after rapamycin stimulation. **b)** Filopodia density in HeLa cells transfected with GA-Split-Cdc42-Q61L was assessed following stimulation with GA_3_-AM for the indicated time. Filopodia density was calculated as the ratio of filopodia number divided by the total cell edge length, using FiloQuant.^1^ Data from over 30 transfected cells per group were combined to generate the violin plots above. Statistical differences were determined using a two-tailed, unpaired student’s *t*-test. ns indicates a p-value of >0.05, * indicates a p-value of <0.05, ** indicates a p-value of <0.01, and *** indicates a p-value of <0.001. On average, a more rapid turn-on is observed in the GA-gated system.

**Figure S6**

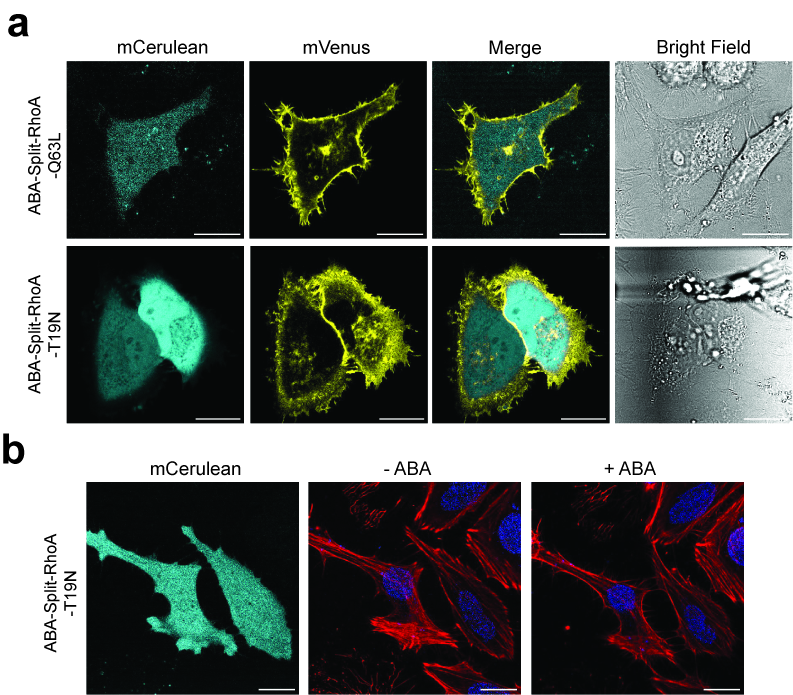

**Figure S6**. Localization of ABA-gated split-RhoA constructs in HeLa cells. **a)** Confocal images of HeLa cells expressing ABA-Split-RhoA-Q63L or ABA-Split-RhoA-T19N constructs. mCerulean fluorescence is localized to the cytosol while mVenus fluorescence is predominantly observed at the plasma membrane. **b)** Confocal images of HeLa cells expressing dominant negative ABA-Split-RhoA-T19N in the absence or presence of 100 µM ABA for 30 min (Red: CellMask Deep Red Actin tracker; Blue: Hoechst stain). In contrast to constitutively active ABA-Split-RhoA-Q63L (**Figure 4b**), cells expressing ABA-Split-RhoA-T19N do not display substantial cell retraction. Scale bar represents 20 µm.

**Materials and Methods**

**Instrumentation and Reagents**

Polymerase chain reaction (PCR) was performed using an Eppendorf thermocycler (05-414-456). Western blots were imaged with a Bio-Rad ChemiDoc XRS+ gel imager (1708265). Protein concentrations were measured using a Synergy H1 hybrid plate reader from Thermo Fisher (11-120-533) for Bradford assays. Mammalian cells were counted with an Invitrogen Countess^TM^ 3 automated cell counter (AMQAX2000). Confocal microscopy was conducted on a Leica STELLARIS 8 system, which features confocal/FLIM/tauSTED capabilities and a tunable white light laser. Cell culture medium was DMEM (Dulbecco’s Modified Eagle Medium, Thermo Fisher, 10569010), supplemented with 10% (v/v) fetal bovine serum,100 U/ml penicillin and 100 g/ml streptomycin. Additional reagents included Gibco^TM^ DPBS (Thermo Fisher, 14-040-133), Gibco^TM^ DMEM (Thermo Fisher, 21-063-029), Gibco^TM^ Opti-MEM^TM^ I Reduced Serum Medium (Thermo Fisher, 11-058-021), and DMSO (Sigma-Aldrich, D8418-100ML). Molecular biology enzymes and kits employed were Platinum Taq DNA polymerase High Fidelity (Thermo Fisher, 11304011), restriction enzymes from New England Biolabs (NEB), T4 DNA ligase (NEB, M0202M), Gibson Assembly Master Mix (NEB, E2611S), and Qiagen plasmid preparation kits (Miniprep Kit, 27104; Maxi Kit, 12162). Chemical inducers such as abscisic acid (TCI Chemicals, A1698-100MG) and gibberellic acid acetoxymethyl ester (GA_3_-AM, Millipore Sigma, SML1959-50MG) were utilized. For cell lysis, protease inhibitor cocktail III (10 µl/ml, Calbiochem, 539134) and phosphatase inhibitor cocktail 1 (10 µl/ml, Sigma, P2825) were used in the lysis buffer. The Bradford protein assay kit from Bio-Rad (5000201) was used for determinations of total protein concentration in lysates. Western blotting employed antibodies from Cell Signaling Technology for Phospho-p44/42 MAPK (Erk1/2) (4377S), P44/42 MAPK (Erk1/2) (9102S), GAPDH (3683S) and HRP-conjugated goat anti-rabbit IgG (7074S). Blots were visualized using Thermo Fisher's SuperSignal^®^ West Dura Extended Duration Chemiluminescent Substrate (34076).

**Cloning**

Constructs used for mammalian transfection were primarily prepared using multiple overlap extension PCR steps followed by restriction enzyme-based cloning. In cases where overlap extension PCR was challenging due to repetitive linker sequences, Gibson assembly was utilized.^2^ The mCerulean and mVenus sequences were sourced from the mCerulean N1 (Addgene, 27795) and mVenus C1 (Addgene, 27794) plasmids, gifts from Steven Vogel at the National Institutes of Health. The N12 and 13C fragments of each small GTPase originated from previously published vectors.^3^ GID1 and GAI_1-92_ were amplified from the pSLQ2812 pPB: CAG-GID1-VPR-IRES-Puro-WPRE PGK-GAI-tagBFP-SpdCas9 plasmid (Addgene, 84240), while PYL and ABI were amplified from the Pslq2817 pPB: CAG-PYL1-VPR-IRES-Puro-WPRE-SV40PA PGK-ABI-tagBFP-SpdCas9 plasmid (Addgene, 84239), both generously provided by Stanley Qi at Stanford University. All plasmids constructed were verified through DNA sequencing. Plasmid information and protein sequences are given in **Tables S1** and **S2**.

**Mammalian cell culture and plasmid transfection**

HeLa cells (ATCC, CCL-2) were cultured in Dulbecco’s modified Eagle’s medium (DMEM) supplemented with 10% (v/v) fetal bovine serum containing 100 U/ml penicillin and 100 g/ml streptomycin. Cells were maintained at 37°C in a 5% CO_2_ humidified incubator. For plasmid transfection, cells were seeded at 1.2 x 10^5^ cells/cm^2^ either on tissue culture treated dishes (Thomas Scientific, 1228K66) for western blot analysis, or on 35 mm poly-d-lysine coated glass-bottom dishes (MatTek Corporation, NC9005934) for confocal microscopy. After 24 hours, the cells had reached ~70-90% confluency and transient transfection was performed following the manufacturer’s instructions (Invitrogen, L3000015).

**Confocal fluorescence imaging**

Confocal images were captured using a Leica STELLARIS 8 confocal/FLIM/tauSTED microscope system equipped with tunable white light lasers and a 37-2 digital temperature controller. The LAS-AF software facilitated image acquisition, with subsequent processing in Fiji (ImageJ). Cells were pre-stained with Hoechst 33342 (Thermo Fisher, H3570) and CellMaskTM Deep Red Actin Tracking Stain (Thermo Fisher, A57245), following the manufacturer’s guidelines, to visualize nuclei and cytoskeletal structures. After staining, cells were rinsed three times with pre-warmed DPBS and incubated in 1.5 ml of DMEM containing 25 mM HEPES (Thermo Fisher, 21-063-029) for imaging. For image acquisition post-CID stimulation, 0.5 ml of DMEM with 4X concentrations of ABA or GA_3_-AM was added dropwise to the dishes using a syringe. Fluorophores were excited with a diode laser at 405 nm or a white light laser spanning 440-790 nm. Nucleus staining was detected with a PMT detector set to capture emission at 430-460 nm following excitation at 405 nm. Actin was visualized with excitation at 652 nm and emission detected between 660-710 nm. mCerulean fluorescence was captured with 458 nm excitation and 468-508 nm emission, while mVenus fluorescence was observed using 514 nm excitation with emission detected at 520-560 nm.

**Lysate preparation and western blot**

Transiently transfected HeLa cells were serum starved overnight in Opti-MEM before being stimulated with GA_3_-AM. Cells were exposed to 10 µM GA_3_-AM in pre-warmed Opti-MEM for the indicated time, followed by rinsing with ice-cold DPBS. Subsequently, cells were lysed using the following lysis buffer: 50 mM Tris-Cl (pH = 7.5 at 25°C), 150 mM NaCl, 50 mM β-glycerophosphate, 10 mM sodium pyrophosphate, 30 mM NaF, 1% Triton X-100, 2 mM EGTA, 100 µM Na_3_VO_4_, 1 mM DTT, protease inhibitor cocktail III (10 µl/ml, Calbiochem, 539134), and phosphatase inhibitor cocktail 1 (10 µl/ml, Sigma, P2825). The lysates were chilled on ice for 15 minutes before centrifugation at 17,000 g for 10 min at 4°C. Protein concentrations were quantified using a Bradford assay (Bio-Rad, 5000201) and normalized for subsequent experiments. For western blot analysis, 5-20 µg of total protein from each lysate was resolved on 12% SDS-PAGE gels and transferred to nitrocellulose membranes. The membranes were then blocked and incubated with the indicated primary antibodies, followed by detection using enhanced chemiluminescence (Thermo Scientific, 34076).

**Data Analysis**

To detect and quantify Cdc42-induced filopodia formation, we utilized the open-source ImageJ plugin, FiloQuant.^1^ This tool enables the extraction of quantitative data, such as the number and length of filopodia and cell edge length.^4^ We evaluated the number of filopodia per cell to assess the efficacy of our GA-split-Cdc42 system on filopodia formation. Data from over 50 transfected cells, collected across multiple cell culture dishes, were aggregated and analyzed. For the time course comparison between GA-split-Cdc42 and Rapa-split-Cdc42 systems, we collected cell images at different time points after CID stimulation (10, 20, and 30 min). Filopodia density, defined as the ratio of filopodia number to total cell edge length, was used to gauge the efficiency of split-Cdc42 systems. For the assessment of ABA-split-RhoA signaling, the cell area was measured using ImageJ to define the region of interest (ROI) for the same transfected cell before and after ABA stimulation. Data for each ABA-split-RhoA construct was gathered from more than 50 cells from multiple cell culture dishes. Statistical analyses were conducted using unpaired, two-tailed student’s *t*-tests (GraphPad Prism 9.5, GraphPad Software).

**Table S1.** Construct description and Addgene IDs.

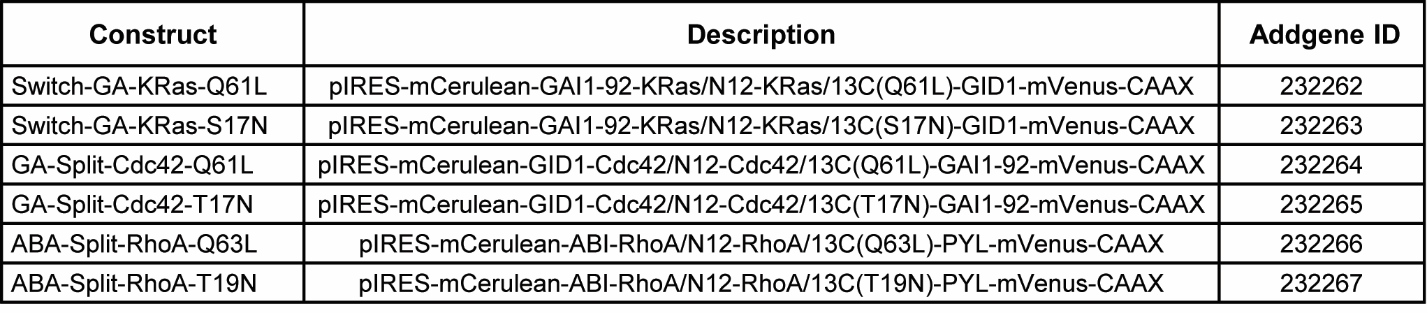

**Table S2.** Amino Acid Sequences for Constructs Used in This Work

1) **pIRES-mCerulean-GAI_1-92_-KRas/N12-KRas/13C(Q61L)-GID1-mVenus-CAAX**

> 61L is underlined.

MVSKGEELFTGVVPILVELDGDVNGHKFSVSGEGEGDATYGKLTLKFICTTGKLPVPWPTLVTTLTWGVQCFARYPDHMKQHDFFKSAMPEGYVQERTIFFKDDGNYKTRAEVKFEGDTLVNRIELKGIDFKEDGNILGHKLEYNAISDNVYITADKQKNGIKANFKIRHNIEDGSVQLADHYQQNTPIGDGPVLLPDNHYLSTQSKLSKDPNEKRDHMVLLEFVTAAGITLGMDELYKGGGSSGGGMKRDHHHHHHQDKKTMMMNEEDDGNGMDELLAVLGYKVRSSEMADVAQKLEQLEVMMSNVQEDDLSQLATETVHYNPAELYTWLDSMLTDLNQISYASRGGGSSGGGELMTEYKLVVVGAG∗---***IRES Region***---MGVGKSALTIQLIQNHFVDEYDPTIEDSYRKQVVIDGETCLLDILDTAG**L**EEYSAMRDQYMRTGEGFLCVFAINNTKSFEDIHHYREQIKRVKDSEDVPMVLVGNKCDLPSRTVDTKQAQDLARSYGIPFIETSAKTRQGVDDAFYTLVREIRKHKEKMSKDGGDPNWELVYTARLQGGGSSGGGQISYASRGMAASDEVNLIESRTVVPLNTWVLISNFKVAYNILRRPDGTFNRHLAEYLDRKVTANANPVDGVFSFDVLIDRRINLLSRVYRPAYADQEQPPSILDLEKPVDGDIVPVILFFHGGSFAHSSANSAIYDTLCRRLVGLCKCVVVSVNYRRAPENPYPCAYDDGWIALNWVNSRSWLKSKKDSKVHIFLAGDSSGGNIAHNVALRAGESGIDVLGNILLNPMFGGNERTESEKSLDGKYFVTVRDRDWYWKAFLPEGEDREHPACNPFSPRGKSLEGVSFPKSLVVVAGLDLIRDWQLAYAEGLKKAGQEVKLMHLEKATVGFYLLPNNNHFHNVMDEISAFVNAECGGGSSGGGVSKGEELFTGVVPILVELDGDVNGHKFSVSGEGEGDATYGKLTLKLICTTGKLPVPWPTLVTTLGYGLQCFARYPDHMKQHDFFKSAMPEGYVQERTIFFKDDGNYKTRAEVKFEGDTLVNRIELKGIDFKEDGNILGHKLEYNYNSHNVYITADKQKNGIKANFKIRHNIEDGGVQLADHYQQNTPIGDGPVLLPDNHYLSYQSKLSKDPNEKRDHMVLLEFVTAAGITLGMDELYKKKKKKKSKTKCVIM∗

2) **pIRES-mCerulean-GAI_1-92_-KRas/N12-KRas/13C(S17N)-GID1-mVenus-CAAX**

> 17N is underlined.

MVSKGEELFTGVVPILVELDGDVNGHKFSVSGEGEGDATYGKLTLKFICTTGKLPVPWPTLVTTLTWGVQCFARYPDHMKQHDFFKSAMPEGYVQERTIFFKDDGNYKTRAEVKFEGDTLVNRIELKGIDFKEDGNILGHKLEYNAISDNVYITADKQKNGIKANFKIRHNIEDGSVQLADHYQQNTPIGDGPVLLPDNHYLSTQSKLSKDPNEKRDHMVLLEFVTAAGITLGMDELYKGGGSSGGGMKRDHHHHHHQDKKTMMMNEEDDGNGMDELLAVLGYKVRSSEMADVAQKLEQLEVMMSNVQEDDLSQLATETVHYNPAELYTWLDSMLTDLNQISYASRGGGSSGGGELMTEYKLVVVGAG∗---***IRES Region***---MGVGK**N**ALTIQLIQNHFVDEYDPTIEDSYRKQVVIDGETCLLDILDTAGQEEYSAMRDQYMRTGEGFLCVFAINNTKSFEDIHHYREQIKRVKDSEDVPMVLVGNKCDLPSRTVDTKQAQDLARSYGIPFIETSAKTRQGVDDAFYTLVREIRKHKEKMSKDGGDPNWELVYTARLQGGGSSGGGQISYASRGMAASDEVNLIESRTVVPLNTWVLISNFKVAYNILRRPDGTFNRHLAEYLDRKVTANANPVDGVFSFDVLIDRRINLLSRVYRPAYADQEQPPSILDLEKPVDGDIVPVILFFHGGSFAHSSANSAIYDTLCRRLVGLCKCVVVSVNYRRAPENPYPCAYDDGWIALNWVNSRSWLKSKKDSKVHIFLAGDSSGGNIAHNVALRAGESGIDVLGNILLNPMFGGNERTESEKSLDGKYFVTVRDRDWYWKAFLPEGEDREHPACNPFSPRGKSLEGVSFPKSLVVVAGLDLIRDWQLAYAEGLKKAGQEVKLMHLEKATVGFYLLPNNNHFHNVMDEISAFVNAECGGGSSGGGVSKGEELFTGVVPILVELDGDVNGHKFSVSGEGEGDATYGKLTLKLICTTGKLPVPWPTLVTTLGYGLQCFARYPDHMKQHDFFKSAMPEGYVQERTIFFKDDGNYKTRAEVKFEGDTLVNRIELKGIDFKEDGNILGHKLEYNYNSHNVYITADKQKNGIKANFKIRHNIEDGGVQLADHYQQNTPIGDGPVLLPDNHYLSYQSKLSKDPNEKRDHMVLLEFVTAAGITLGMDELYKKKKKKKSKTKCVIM∗

3) **pIRES-mCerulean-GID1-Cdc42/N12-Cdc42/13C(Q61L)-GAI_1-92_-mVenus-CAAX**

> 61L is underlined.

MVSKGEELFTGVVPILVELDGDVNGHKFSVSGEGEGDATYGKLTLKFICTTGKLPVPWPTLVTTLTWGVQCFARYPDHMKQHDFFKSAMPEGYVQERTIFFKDDGNYKTRAEVKFEGDTLVNRIELKGIDFKEDGNILGHKLEYNAISDNVYITADKQKNGIKANFKIRHNIEDGSVQLADHYQQNTPIGDGPVLLPDNHYLSTQSKLSKDPNEKRDHMVLLEFVTAAGITLGMDELYKGGGSSGGGMAASDEVNLIESRTVVPLNTWVLISNFKVAYNILRRPDGTFNRHLAEYLDRKVTANANPVDGVFSFDVLIDRRINLLSRVYRPAYADQEQPPSILDLEKPVDGDIVPVILFFHGGSFAHSSANSAIYDTLCRRLVGLCKCVVVSVNYRRAPENPYPCAYDDGWIALNWVNSRSWLKSKKDSKVHIFLAGDSSGGNIAHNVALRAGESGIDVLGNILLNPMFGGNERTESEKSLDGKYFVTVRDRDWYWKAFLPEGEDREHPACNPFSPRGKSLEGVSFPKSLVVVAGLDLIRDWQLAYAEGLKKAGQEVKLMHLEKATVGFYLLPNNNHFHNVMDEISAFVNAECQISYASRGGGSSGGGELQTIKCVVVGDGA∗---***IRES Region***---MGVGKTCLLISYTTNKFPSEYVPTVFDNYAVTVMIGGEPYTLGLFDTAG**L**EDYDRLRPLSYPQTDVFLVCFSVVSPSSFENVKEKWVPEITHHCPKTPFLLVGTQIDLRDDPSTIEKLAKNKQKPITPETAEKLARDLKAVKYVECSALTQRGLKNVFDEAILAALEPPETQPGDPNWELVYTARLQGGGSSGGGQISYASRGMKRDHHHHHHQDKKTMMMNEEDDGNGMDELLAVLGYKVRSSEMADVAQKLEQLEVMMSNVQEDDLSQLATETVHYNPAELYTWLDSMLTDLNGGGSSGGGVSKGEELFTGVVPILVELDGDVNGHKFSVSGEGEGDATYGKLTLKLICTTGKLPVPWPTLVTTLGYGLQCFARYPDHMKQHDFFKSAMPEGYVQERTIFFKDDGNYKTRAEVKFEGDTLVNRIELKGIDFKEDGNILGHKLEYNYNSHNVYITADKQKNGIKANFKIRHNIEDGGVQLADHYQQNTPIGDGPVLLPDNHYLSYQSKLSKDPNEKRDHMVLLEFVTAAGITLGMDELYKKKKKKKSKTKCVIM∗

4) **pIRES-mCerulean-GID1-Cdc42/N12-Cdc42/13C(T17N)-GAI_1-92_-mVenus-CAAX**

> 17N is underlined.

MVSKGEELFTGVVPILVELDGDVNGHKFSVSGEGEGDATYGKLTLKFICTTGKLPVPWPTLVTTLTWGVQCFARYPDHMKQHDFFKSAMPEGYVQERTIFFKDDGNYKTRAEVKFEGDTLVNRIELKGIDFKEDGNILGHKLEYNAISDNVYITADKQKNGIKANFKIRHNIEDGSVQLADHYQQNTPIGDGPVLLPDNHYLSTQSKLSKDPNEKRDHMVLLEFVTAAGITLGMDELYKGGGSSGGGMAASDEVNLIESRTVVPLNTWVLISNFKVAYNILRRPDGTFNRHLAEYLDRKVTANANPVDGVFSFDVLIDRRINLLSRVYRPAYADQEQPPSILDLEKPVDGDIVPVILFFHGGSFAHSSANSAIYDTLCRRLVGLCKCVVVSVNYRRAPENPYPCAYDDGWIALNWVNSRSWLKSKKDSKVHIFLAGDSSGGNIAHNVALRAGESGIDVLGNILLNPMFGGNERTESEKSLDGKYFVTVRDRDWYWKAFLPEGEDREHPACNPFSPRGKSLEGVSFPKSLVVVAGLDLIRDWQLAYAEGLKKAGQEVKLMHLEKATVGFYLLPNNNHFHNVMDEISAFVNAECQISYASRGGGSSGGGELQTIKCVVVGDGA∗---***IRES Region***---MGVGK**N**CLLISYTTNKFPSEYVPTVFDNYAVTVMIGGEPYTLGLFDTAGQEDYDRLRPLSYPQTDVFLVCFSVVSPSSFENVKEKWVPEITHHCPKTPFLLVGTQIDLRDDPSTIEKLAKNKQKPITPETAEKLARDLKAVKYVECSALTQRGLKNVFDEAILAALEPPETQPGDPNWELVYTARLQGGGSSGGGQISYASRGMKRDHHHHHHQDKKTMMMNEEDDGNGMDELLAVLGYKVRSSEMADVAQKLEQLEVMMSNVQEDDLSQLATETVHYNPAELYTWLDSMLTDLNGGGSSGGGVSKGEELFTGVVPILVELDGDVNGHKFSVSGEGEGDATYGKLTLKLICTTGKLPVPWPTLVTTLGYGLQCFARYPDHMKQHDFFKSAMPEGYVQERTIFFKDDGNYKTRAEVKFEGDTLVNRIELKGIDFKEDGNILGHKLEYNYNSHNVYITADKQKNGIKANFKIRHNIEDGGVQLADHYQQNTPIGDGPVLLPDNHYLSYQSKLSKDPNEKRDHMVLLEFVTAAGITLGMDELYKKKKKKKSKTKCVIM∗

5) **pIRES-mCerulean-ABI-RhoA/N12-RhoA/13C(Q63L)-PYL-mVenus-CAAX**

> 63L is underlined.

MVSKGEELFTGVVPILVELDGDVNGHKFSVSGEGEGDATYGKLTLKFICTTGKLPVPWPTLVTTLTWGVQCFARYPDHMKQHDFFKSAMPEGYVQERTIFFKDDGNYKTRAEVKFEGDTLVNRIELKGIDFKEDGNILGHKLEYNAISDNVYITADKQKNGIKANFKIRHNIEDGSVQLADHYQQNTPIGDGPVLLPDNHYLSTQSKLSKDPNEKRDHMVLLEFVTAAGITLGMDELYKGGGSSGGGVPLYGFTSICGRRPEMEAAVSTIPRFLQSSSGSMLDGRFDPQSAAHFFGVYDGHGGSQVANYCRERMHLALAEEIAKEKPMLCDGDTWLEKWKKALFNSFLRVDSEIESVAPETVGSTSVVAVVFPSHIFVANCGDSRAVLCRGKTALPLSVDHKPDREDEAARIEAAGGKVIQWNGARVFGVLAMSRSIGDRYLKPSIIPDPEVTAVKRVKEDDCLILASDGVWDVMTDEEACEMARKRILLWHKKNAVAGDASLLADERRKEGKDPAAMSAAEYLSKLAIQRGSKDNISVVVVDLKQISYASRGGGSSGGGELAAIRKKLVIVGDGA∗---***IRES Region***---MCGKTCLLIVFSKDQFPEVYVPTVFENYVADIEVDGKQVELALWDTAG**L**EDYDRLRPLSYPDTDVILMCFSIDSPDSLENIPEKWTPEVKHFCPNVPIILVGNKKDLRNDEHTRRELAKMKQEPVKPEEGRDMANRIGAFGYMECSAKTKDGVREVFEMATRAALQAGDPNWELVYTARLQGGGSSGGGQISYASRGTQDEFTQLSQSIAEFHTYQLGNGRCSSLLAQRIHAPPETVWSVVRRFDRPQIYKHFIKSCNVSEDFEMRVGCTRDVNVISGLPANTSRERLDLLDDDRRVTGFSITGGEHRLRNYKSVTTVHRFEKEEEEERIWTVVLESYVVDVPEGNSEEDTRLFADTVIRLNLQKLASITEAMNGGGSSGGGVSKGEELFTGVVPILVELDGDVNGHKFSVSGEGEGDATYGKLTLKLICTTGKLPVPWPTLVTTLGYGLQCFARYPDHMKQHDFFKSAMPEGYVQERTIFFKDDGNYKTRAEVKFEGDTLVNRIELKGIDFKEDGNILGHKLEYNYNSHNVYITADKQKNGIKANFKIRHNIEDGGVQLADHYQQNTPIGDGPVLLPDNHYLSYQSKLSKDPNEKRDHMVLLEFVTAAGITLGMDELYKKKKKKKSKTKCVIM∗

6) **pIRES-mCerulean-ABI-RhoA/N12-RhoA/13C(T19N)-PYL-mVenus-CAAX**

> 19N is underlined.

MVSKGEELFTGVVPILVELDGDVNGHKFSVSGEGEGDATYGKLTLKFICTTGKLPVPWPTLVTTLTWGVQCFARYPDHMKQHDFFKSAMPEGYVQERTIFFKDDGNYKTRAEVKFEGDTLVNRIELKGIDFKEDGNILGHKLEYNAISDNVYITADKQKNGIKANFKIRHNIEDGSVQLADHYQQNTPIGDGPVLLPDNHYLSTQSKLSKDPNEKRDHMVLLEFVTAAGITLGMDELYKGGGSSGGGVPLYGFTSICGRRPEMEAAVSTIPRFLQSSSGSMLDGRFDPQSAAHFFGVYDGHGGSQVANYCRERMHLALAEEIAKEKPMLCDGDTWLEKWKKALFNSFLRVDSEIESVAPETVGSTSVVAVVFPSHIFVANCGDSRAVLCRGKTALPLSVDHKPDREDEAARIEAAGGKVIQWNGARVFGVLAMSRSIGDRYLKPSIIPDPEVTAVKRVKEDDCLILASDGVWDVMTDEEACEMARKRILLWHKKNAVAGDASLLADERRKEGKDPAAMSAAEYLSKLAIQRGSKDNISVVVVDLKQISYASRGGGSSGGGELAAIRKKLVIVGDGA∗---***IRES Region***---MCGK**N**CLLIVFSKDQFPEVYVPTVFENYVADIEVDGKQVELALWDTAGQEDYDRLRPLSYPDTDVILMCFSIDSPDSLENIPEKWTPEVKHFCPNVPIILVGNKKDLRNDEHTRRELAKMKQEPVKPEEGRDMANRIGAFGYMECSAKTKDGVREVFEMATRAALQAGDPNWELVYTARLQGGGSSGGGQISYASRGTQDEFTQLSQSIAEFHTYQLGNGRCSSLLAQRIHAPPETVWSVVRRFDRPQIYKHFIKSCNVSEDFEMRVGCTRDVNVISGLPANTSRERLDLLDDDRRVTGFSITGGEHRLRNYKSVTTVHRFEKEEEEERIWTVVLESYVVDVPEGNSEEDTRLFADTVIRLNLQKLASITEAMNGGGSSGGGVSKGEELFTGVVPILVELDGDVNGHKFSVSGEGEGDATYGKLTLKLICTTGKLPVPWPTLVTTLGYGLQCFARYPDHMKQHDFFKSAMPEGYVQERTIFFKDDGNYKTRAEVKFEGDTLVNRIELKGIDFKEDGNILGHKLEYNYNSHNVYITADKQKNGIKANFKIRHNIEDGGVQLADHYQQNTPIGDGPVLLPDNHYLSYQSKLSKDPNEKRDHMVLLEFVTAAGITLGMDELYKKKKKKKSKTKCVIM∗
